# SnpReady for Rice (SR4R) Database

**DOI:** 10.1101/2020.01.11.902999

**Authors:** Jun Yan, Dong Zou, Chen Li, Zhang Zhang, Shuhui Song, Xiangfeng Wang

**Affiliations:** Department of Crop Genomics and Bioinformatics, College of Agronomy and Biotechnology, China Agricultural University, Beijing 100094, China; National Genomics Data Center & BIG Data Center & CAS Key Laboratory of Genome Sciences and Information, Beijing Institute of Genomics, Chinese Academy of Sciences, Bejing 100101, China; Rice Research Institute, Guangdong Academy of Agricultural Sciences, Guangzhou 510640, China

**Keywords:** Rice, SNP, Database, Hapmap

## Abstract

The information commons for rice (IC4R) database is a collection of ∼18 million SNPs (single nucleotide polymorphisms) identified by the resequencing of 5,152 rice accessions. Although IC4R offers ultra-high density rice variation map, these raw SNPs are not readily usable for the public. To satisfy different research utilizations of SNPs for population genetics, evolutionary analysis, association studies and genomic breeding in rice, the raw genotypic data of the 18 million SNPs were processed by unified bioinformatics pipelines. The outcomes were used to develop a daughter database of IC4R – SnpReady for Rice (SR4R). The SR4R presents four reference SNP panels, including 2,097,405 hapmapSNPs after data filtration and genotype imputation, 156,502 tagSNPs selected from linkage disequilibrium (LD)-based redundancy removal, 1,180 fixedSNPs selected from genes exhibiting selective sweep signatures, and 38 barcodeSNPs selected from DNA fingerprinting simulation. SR4R thus offers a highly efficient rice variation map that combines reduced SNP redundancy with extensive data describing the genetic diversity of rice populations. In addition, SR4R provides rice researchers with a web-interface that enables them to browse all four SNP panels, use online toolkits, and retrieve the original data and scripts for a variety of population genetics analyses on local computers. The SR4R is freely available to academic users at http://sr4r.ic4r.org/.

## Introduction

*Oryza sativa*, or rice, was the first crop genome to be sequenced. In the past decade, thousands of rice accessions in the germplasm banks worldwide have been genotyped [1] and numerous rice variation databases have been constructed. One of these databases is the rice variation database (RVD), a daughter database of the Information Commons for Rice consortium (IC4R) [2]. RVD is a collection of over eighteen million SNPs (single nucleotide polymorphisms) identified from 5,152 rice accessions based on whole-genome resequencing data, and offers an ultra-high-density rice variation map – about one SNP per twenty bases on average. The information contained in this high volume of raw SNPs is not ready for use until it has been processed to remove low-quality SNPs, such as those with missing/low frequency genotypes, or redundant SNPs identified due to linkage disequilibrium (LD). In addition, different types of research require different magnitudes of SNPs to ensure efficient computing and accurate results; for example, the requirements are different for evolutionary studies using comparative genomics and pan-genome analysis, gene mapping by quantitative trait loci (QTL), genome-wide association study analysis (GWAS), molecular breeding by marker-assisted selection (MAS) and genomic selection (GS), and variety protection by DNA fingerprint barcoding.

Construction of a reference haplotype map (HapMap) to represent the maximal population diversity for a species is the first step. The ∼18 million raw SNPs in RVD provide an initial variation dataset to generate a reference HapMap for rice. According to international human HapMap database, which contains over 3.1 million high-quality SNPs, a density of one SNP per 100 bases is sufficient for performing genotype imputation, GWAS analysis and mapping of causal variations [3]. Because the genome size of rice is ∼ 400 Mbp, about two million high-quality SNPs may offer an ideal density of one SNP per 200 bases. Such density of a reference rice HapMap is especially useful for molecular breeders to perform genotype imputation to supplement missing genotypes or increase SNP density, as low-density genotyping platforms are mostly used in rice to lower genotyping expense.

For population genetics studies in which thousands of individual samples are assessed, the millions of SNPs in an entire HapMap are excessive. The redundant SNPs in a HapMap extensively increase computing costs, and may also reduce the accuracy of results. To circumvent these challenges, a subgroup of SNPs whose genotypes significantly correlate with other SNPs in the same linkage disequilibrium (LD) region are selected; these are known as tagging SNPs. The number of tagging SNPs may vary between species and populations, depending upon the lengths of LD regions in each group [4]. Based on the data in RVD, LD length in rice ranges from 100 to 500 Kb; thus 100,000 SNPs, which yields a density of one tagging SNP per 3 to 5 Kbp, is sufficient for various genetic diversity analysis.

The expense of genotyping is an important factor to consider in crop molecular breeding, as molecular breeding typically requires the rapid genotyping of thousands of samples, often within days or even hours. Therefore, low SNP density genotyping technologies, such as SNP chip or KASP-based platforms are usually preferred by industrial seed companies; these methods offer great flexibility by combining the rapid identification of low numbers of SNPs (several to a few dozen) with the ability to multiplex hundreds to thousands of DNA samples. However, these methods lack precision.

Modern breeding methods demand the efficiency and stability of a highly concise marker panel containing ∼1K SNPs. SNPs used to select plants for breeding typically occur in genes or genomic regions that are associated with agronomic traits believed to be subject to selective pressures [5]. Genes with variations exhibiting selectively fixed signatures can be identified based on the *θπ* and *Fst* values computed by selective sweep analysis [6]. This magnitude of SNPs is suitable for synthesis on low-density SNP chips, which are then used for conducting certain types of molecular analysis, such as marker-assisted selection, seed purity or heterozygosity testing, genetic component analysis, and subpopulation classification. For intellectual protection of commercial rice varieties, DNA fingerprinting typically uses only 12 to 36 SNPs, to generate a combination of barcodes with maximal resolution to distinguish commercial varieties in the seed industry or germplasm accessions in gene banks. Simulation of all possible combinations of a set of candidate SNPs have to be tested in a large germplasm population to ensure the maximal resolution with fewest markers, such as the MinimalMarker algorithm [7].

To enhance the ability of researchers to effectively use the RVD in IC4R, we developed a daughter database we have called SnpReady for Rice, or SR4R. SR4R enables researcher to readily retrieve SNPs that are relevant to their own research, thus saving time and computational resources. In SR4R, the ∼18 million SNPs have been divided into four categories: hapmapSNPs, tagSNPs, fixedSNPs, and barcodeSNPs (**Figure 1**). SR4R allows users to browse the related information associated with each SNP panel, and also to download each set of genotype files for local use. SR4R also offers 18 bioinformatics tools and pipeline scripts, enabling users to locally run the tools to perform genotype imputation, basic statistical analysis, genotype file format conversion, SNPs filtration and extraction, population structure analysis, genetic diversity analysis, rice subpopulation classification, DNA fingerprinting analysis, and other additional functions.

**Figure 1.**
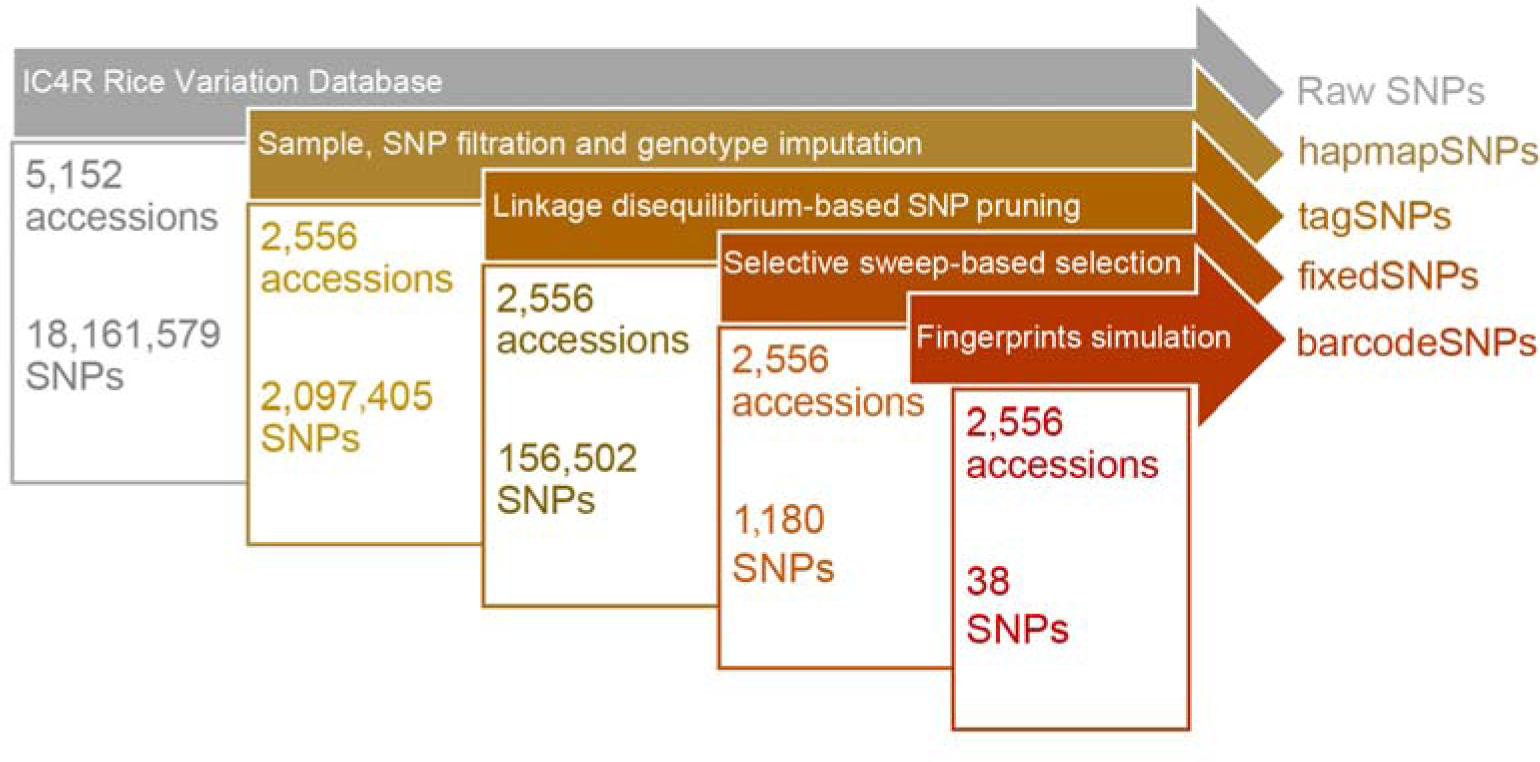
An overview of the four SNP panels of the SR4R database. The flow chart describes procedures on how the four SNP panels were generated.

### Database contents and analytical modules

#### The hapmapSNP panel

The IC4R rice variation database (RVD; http://variation.ic4r.org/) is a collection of over 18 million SNPs with related annotation information, identified from previously published whole genome resequencing of 5,152 rice accessions [2]. Such a high-density rice variation map, which identifies an average of one SNP per twenty bases, offers the possibility of generating a high-density HapMap for the rice research community; creating such a HapMap was the first step in creating the SnpReady for Rice (SR4R) Database described here.

To ensure the quality of HapMap, we performed an initial filtration of samples and SNPs on the raw dataset of 5,152 accessions (**Materials and Methods**). First, a total of 2,556 accessions with genotype missing rate less than 20% were selected; each selected accession has been documented with explicit subpopulation classification and origins (**Table S1**). Then, SNPs with genotype missing rate ≥ 0.1 and minor allele frequency (MAF) ≤ 0.05 were removed. Genotype imputation on the resulting 2,883,623 SNPs in the selected 2,556 accessions yielded a high-quality HapMap containing 2,097,405 SNPs without any missing genotypes using the software Beagle [8]. These 2,097,405 SNPs were regarded as the hapmapSNP panel, and were used as the initial dataset for generating the other three SNP panels (**Figure 2A and 2D**).

**Figure 2.**
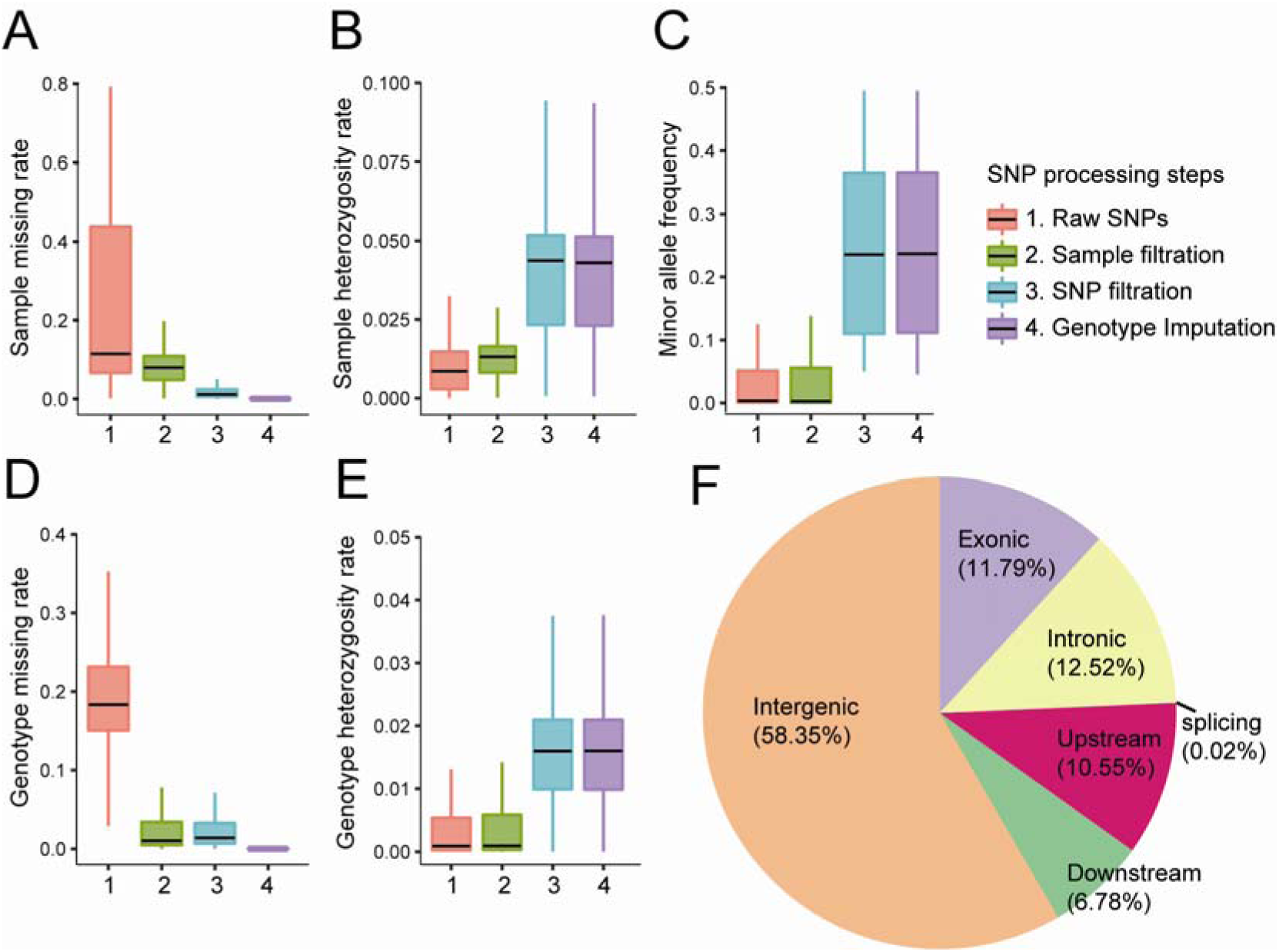
Basic statistics of the rice hapmapSNPs after four steps of genotype processing. After four steps of genotype processing, a series of basic statistical analyses was performed at each step to exhibit the characteristics of the SNPs, including: **A**. statistics of individual missing rate, **B**. statistics of individual heterozygote rate, **C**. statistics of minor allele frequency, **D**. statistics of site missing rate, and **E**. statistics of site heterozygote rate. **F.** After ARNOVAR analysis to annotated the hapmapSNPs, distribution of the hapmapSNPs in different genomic regions were analyzed.

The generated reference HapMap of rice has an average density of five SNPs per Kb with a heterozygosity rate of 3.75% (**Figure 2B and 2D**). Genome-wide distribution statistics showed that 58.4% of the hapmapSNPs present in the intergenic regions, 12.5% in the intronic regions, 11.8% in the exonic regions, 0.02% on the splicing sites, and 10.6% and 6.8% hapmapSNPs located in the upstream and downstream regions (1Kb away from transcription start site or transcription end site) of a gene territory (**Figure 2F**). The 2,097,405 hapmapSNPs with genotypes of 2,556 accessions are available to download, enabling users to perform genotype imputation on local genotype data to increase the density of SNPs generated from low-density genotyping platform.

#### The tagSNP panel

High SNP density is usually beneficial to precise mapping of trait-related genes with GWAS analysis, but is not suitable for population genetic analysis because SNP redundancy may add unnecessary computation costs and introduce bias to the results [9]. Since SNPs within the same LD region possess correlated genotypes forming one haplotype block, a representative SNP is usually selected as a tag to solve the redundancy issue. We adopted an LD-based SNP pruning procedure to infer haplotype tagging SNPs (tagSNPs) from the hapmapSNPs (**Materials and Methods**). As a result, 156,502 tagSNPs were identified **(Figure 1).** To verify whether the tagSNP panel properly represents the genetic diversity of the population, phylogenetic analysis using the 156,502 tagSNPs was performed on the 2,556 rice accessions which were explicitly documented with subpopulation classification and origins. As shown in **Figure 3A**, the resulting phylogenetic tree clearly exhibited six major clades representing the five cultivated rice subpopulations and one wild rice subpopulation. The five cultivated rice subpopulations include *indica* rice (*Ind* for short) containing 1,655 accessions, *Aus* rice (*Aus*) containing 182 accessions, *Aromatic* (*Aro*) rice containing 56 accessions, tropical *japonica* rice (*TrJ*) containing 318 accessions, and temperate *japonica* rice (*Tej*) containing 327 accessions, whilst the wild rice subpopulation contains 18 *O. rufipogon* (*Oru*) accessions. In addition, PCA-based (**Figure 3B**) and admixture-based (**Figure 3C**) analyses showed the same pattern, with the subpopulation classification as the phylogenetic tree indicated. For population admixture structure analysis, a predefined parameter of “K value” was used to mandatorily estimate the number of ancestral subpopulation and uses different colours for each K value to represent the number of subpopulations. Because the optimal number of ancestral subpopulation is usually unknown, a common way is to use a series of K value to estimate the optimal K parameter. It is worth noting that the *japonica, indica* and *Aus* subpopulations were explicitly separated when K was set to 3, while the six subpopulations were clearly separated until the K value was set to 8. In addition, between K=4 to 7, the *indica* subpopulation showed clear structure divided into six groups (*indica* g1 to g6) as indicated by both PCA and admixture analysis (**Figure 3D and Figure S1**). The genetic structures of the six rice subpopulations and the six *indica* subgroups are consistent with multiple previous reports [10].

**Figure 3.**
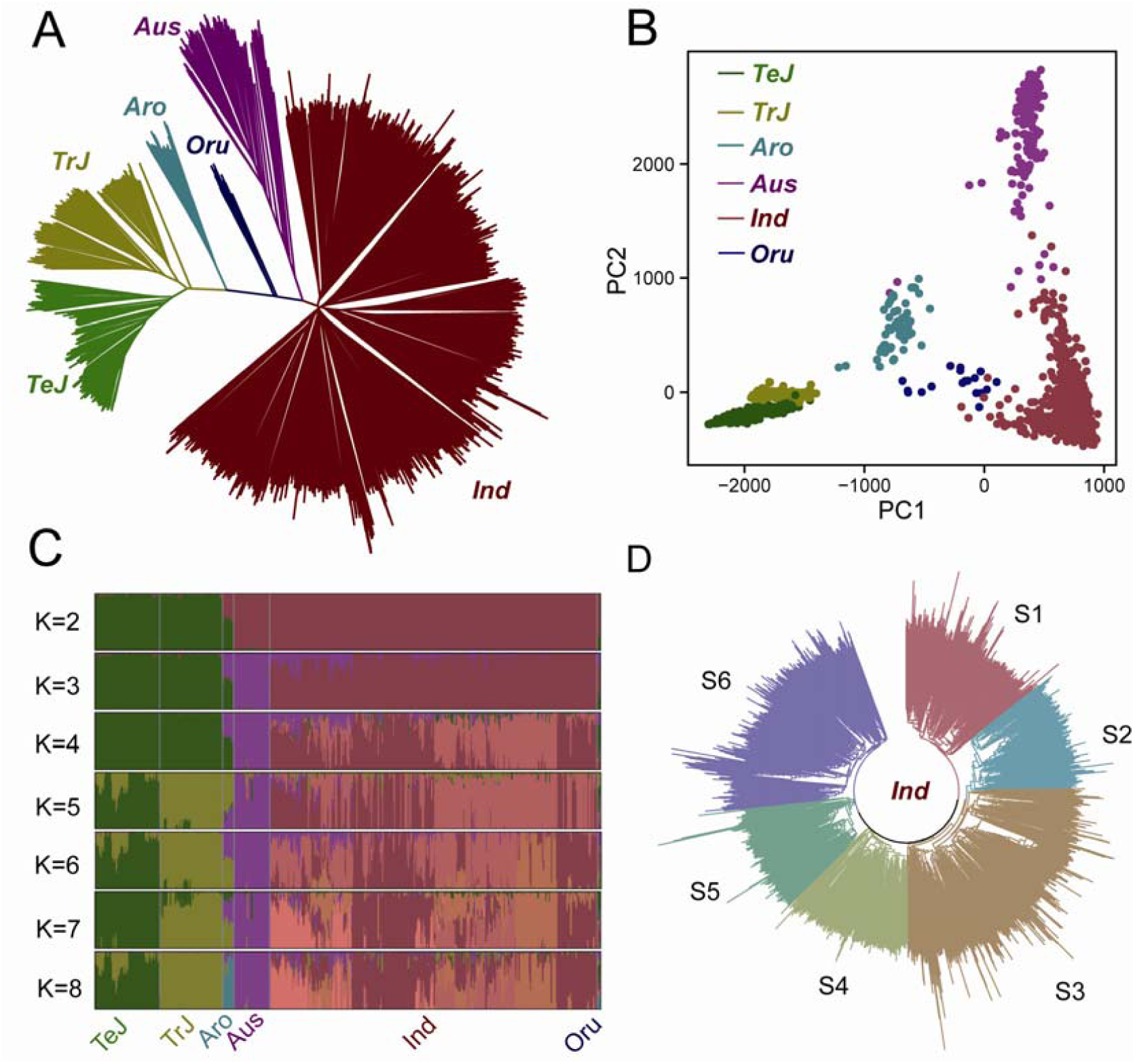
Population structure analysis of the 2,556 rice accessions using tagSNPs. To test whether the 150K tagSNPs can generate the population structures consistent with previous reports, we performed a series of population structure analyses to generate: **A**. the phylogenetic tree, **B.** the PCA map, **C.** the admixture structure of 2,556 rice accessions, **D.** the phylogenetic tree of the six subgroups of *indica* rice. The tagSNPs effectively and accurately classified the 2,556 rice accessions to corresponding populations.

#### Genetic diversity analysis with the tagSNP panel

The tagSNP panel represents a subset of the hapmapSNPs after approximately 92.5% of the genetic redundancy was removed (**Figure 1**). To test the effectiveness of the 156,502 tagSNPs, we performed another series of standard genetic diversity analyses and examined whether the results agreed with previously reported conclusions. First, we found that the count of homozygous SNPs and the heterozygosity rate of the accessions in the six subpopulations showed opposite trends: while the accessions in the *TeJ* subpopulation had the highest count of homozygous SNPs and lowest heterozygous rate, the accessions in the *indica* subpopulation had the lowest count of homozygosity SNPs and highest homozygosity rate (**Figure 4A and 4B**). The IBS (identity by state) analysis is a commonly used method to measure the similarity of alleles in a designated population, which may reflect the genetic diversity of the whole population and subpopulations. Comparison of the IBS values among different subpopulations may help understand the degree of genetic differentiation in different subpopulations. In order to validate whether the IBS results generated from the tagSNPs are consistent with the previous reports regarding the genetic diversity in different subpopulations, pairwise computation of the IBS values between each pair of accessions within the same subpopulation was performed, and the results showed that temperate *japonica* rice has the highest IBS values, while the *indica* rice has the lowest (**Figure 4C**). In addition, runs of homozygosity (ROH) analysis indicated that the temperate *japonica* rice has the most and longest ROH regions, while the *indica* rice has the least and shortest ROH regions (**Figure 4D**). This pattern agreed with the result from LD decay analysis showing that temperate *japonica* rice has the slowest LD decay rate while the *indica* rice has fastest rate (**Figure 4E**). Computations of *θπ* and *Fst* are commonly used methods to measure genetic diversity within population and between population, respectively (**Materials and Methods**). The within-subpopulation diversities of the six rice subpopulations are *Oru (θπ*=0.218), *Ind* (*θπ*=0.216), *Aus* (*θπ*=0.182), *Aro* (*θπ*=0.145), *TrJ* (*θπ*=0.116) and *TeJ* (*θπ*=0.068). Using the wild rice subpopulation as reference, the genetic distances of the five types of cultivated rice between wild rice are *TeJ* (*Fst*=0.476), *TrJ* (*Fst*=0.419), *Aus* (*Fst*=0.299), *Ind* (*Fst*=0.266) and *Aro* (*Fst*=0.241), suggesting the highest domestication level of *japonica* rice compared to other rice (**Figure 4F and 4G**). The collective results from multiple angles of standard genetic diversity analyses were consistent with previous reports that *indica* rice has a more complicated genetic composition and origin compared to the other five subpopulations [11].

**Figure 4.**
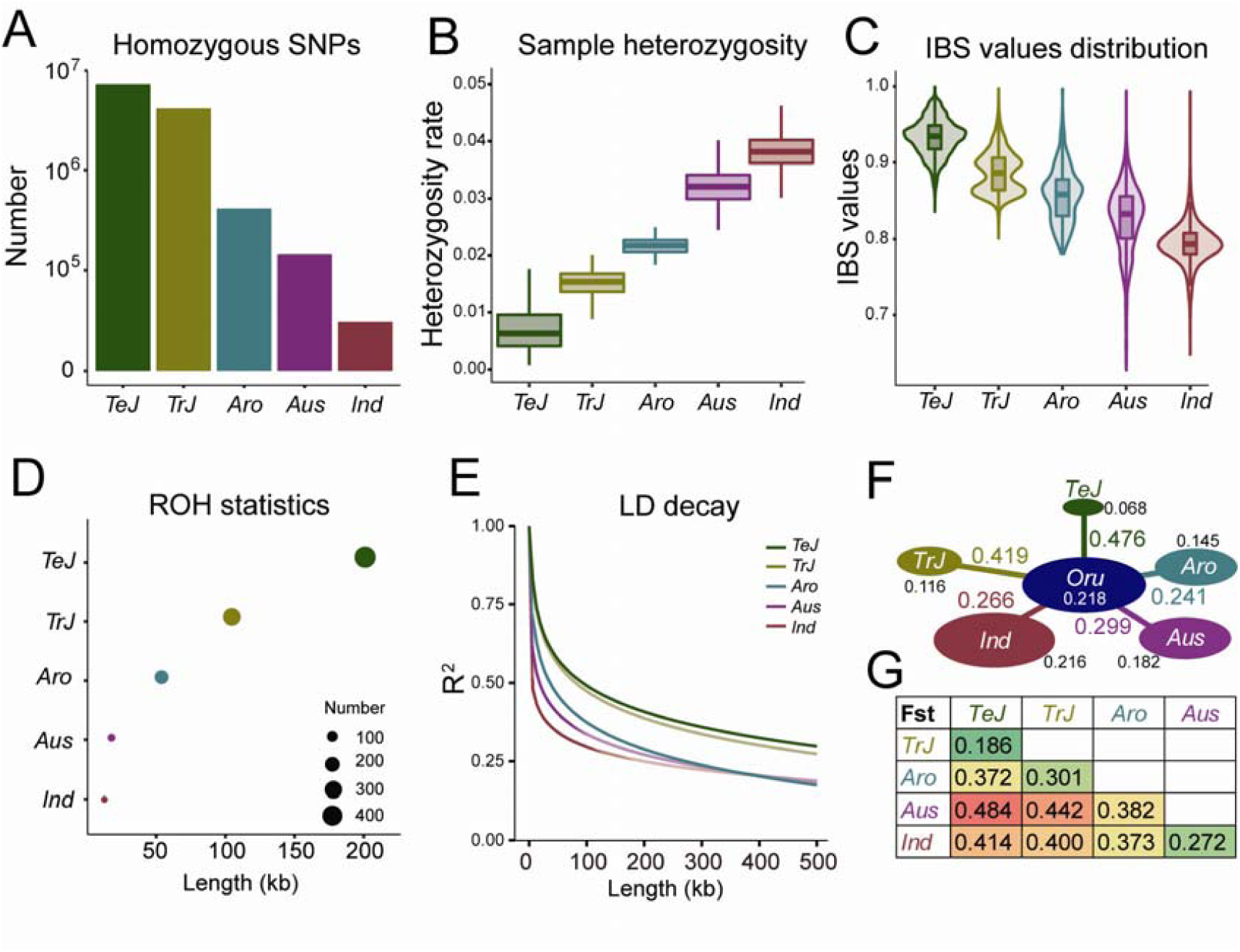
Genetic diversity analysis of rice accessions using tagSNPs. The 150K tagSNPs were used in a series of population genetic analysis to show the effectiveness of tagSNPs including: **A**. statistics of homozygous SNPs, **B.** statistics of individual heterozygosity, **C**. pairwise IBS values distribution, **D.** statistics of ROH regions, **E**. LD decay analysis, in the five major rice subpopulations. **F**. Genetic diversity (*θπ*) and population differentiation (*Fst*) between cultivated and wild subpopulations. **G**. Population differentiation (*Fst*) of cultivated subpopulations.

#### Genomic selection analysis with the tagSNP panel

Genomic selection (GS) has been widely used in industrial animal and crop breeding programs [12]. GS is essentially a best linear unbiased prediction (BLUP) model that is first trained with known genotypes and phenotypes of reference population individuals, usually accounting for 20% to 50% of a breeding population, and then used to predict the unknown phenotypes of the remaining genotyped individuals (the candidate population). The predicted phenotypes, known as the genomic estimated breeding values (GEBV), are ranked from high to low, and can be used to assist in deciding upon a hybridization plan. Although GS may significantly shorten the breeding cycle, the cost for genotyping has been a vital factor because the GS model has to take genome-wide SNP markers as input, especially from crop breeding in which thousands to hundreds of thousands of individuals need to be genotyped. In order to lower genotyping cost, compilation of a set of thousands of SNPs that may best represent the overall genetic backgrounds of a breeding population is of great importance.

Because the 156,502 tagSNPs category is a high-quality marker set with most redundancy removed while preserving maximal genetic diversity, it may be considered as a marker pool for selecting high-efficiency SNPs for genomic selection. To test the effectiveness, we analysed a previously published dataset containing 414 rice parental lines with non-missing genotypes of 29,434 SNPs profiled by the 44K rice SNP chip, and nine phenotype traits (flowering time, panicle fertility, seed width, seed volume, seed surface area, plant height, flag leaf length, flag leaf width, and florets per panicle). The GS model was obtained from the ridge regression best linear unbiased prediction (rrBLUP) algorithm [13], and prediction accuracy was evaluated with Pearson correlation between observed and predicted traits by five-fold cross validation. The evaluation was performed using five different SNP combinations: Set-1, the original 29,434 SNPs on the 44K chip; Set-2, the 1,090 SNPs overlapped between the 156,502 tagSNPs and 29,434 SNPs; Set-3, the 1,090 SNPs randomly selected from the 29,434 SNPs; Set-4, the 1,090 SNPs evenly distributed in the genome (350 Kb per SNP) selected from the 29,434 SNPs; and Set-5, the 1,090 consecutive SNPs localized within a randomly selected genomic region from the 29,434 SNPs. Then the rrBLUP prediction was performed on the nine phenotypes using the five sets of SNPs to compare prediction accuracies (**Figure 5**). Although prediction accuracies greatly varied ranging from 0.23 to 0.90 among the nine traits due to different heritability of each trait, the trend of the five SNP sets within the same trait was generally consistent. Except for the trait of panicle fertility in which the Set-2 SNPs exhibited the highest prediction accuracy, the full 29,434 SNPs showed the highest prediction accuracy for the other eight traits followed by the 1,090 tagSNPs in the second position. We further performed pairwise student’s t-test for Pearson correlations of the selected 1,090 tagSNPs set (Set-2) and other four sets, the result shows that the selected 1,090 tagSNPs set significantly outperform other randomly selected SNP set for most traits (**Figure S2**). These results indicate that selection of about one thousand tagSNPs from the tagSNP pool might be a feasible option to lower genotyping budget; for example, these SNPs could inform the synthesis of a new low-density SNP chip rather than using high-density SNP chip.

**Figure 5.**
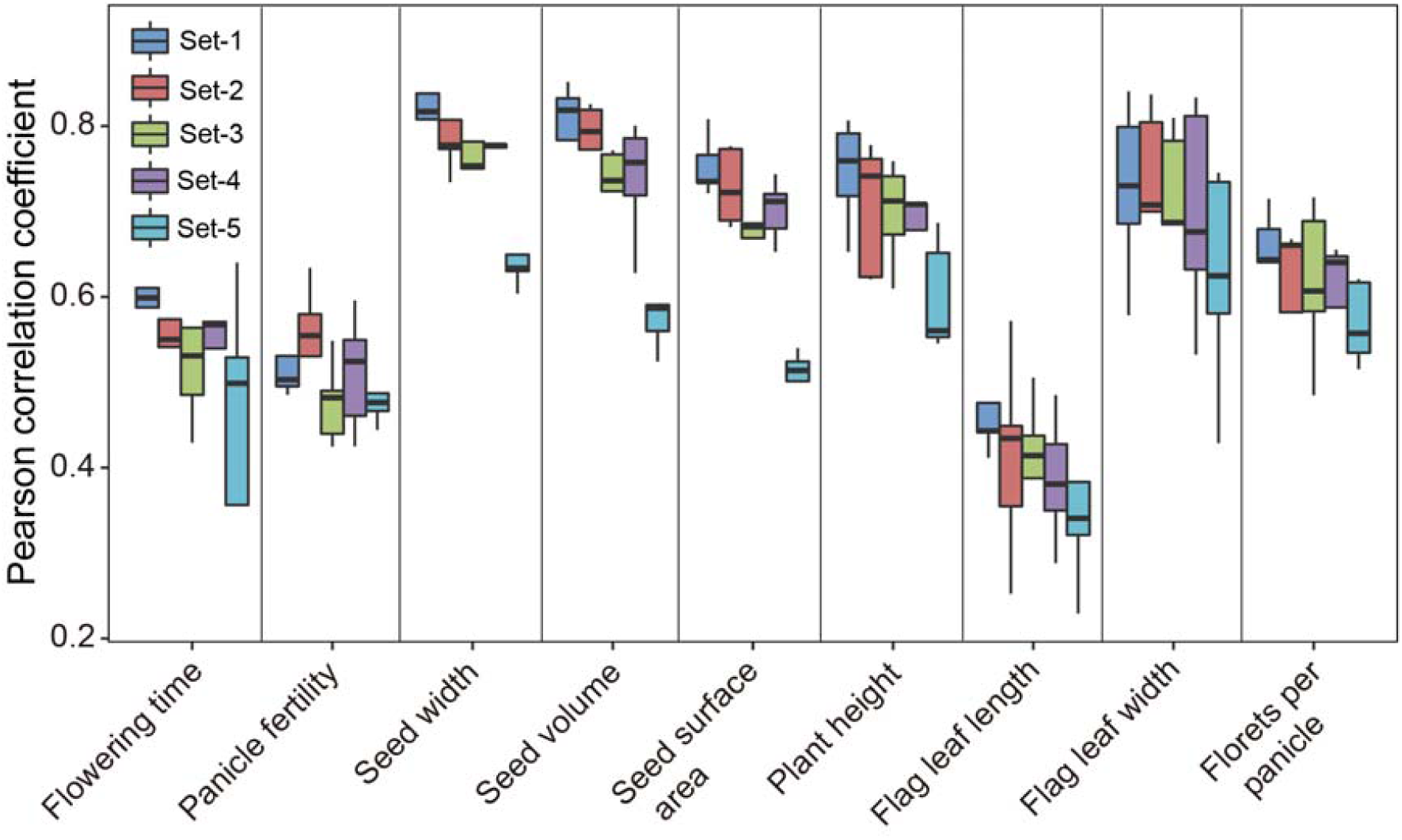
Genomic selection-based phenotype prediction using tagSNPs. Nine agronomical phenotypes were predicted based on rrBLUP models to evaluate the effectiveness of tagSNPs. Five sets of SNPs with equal amounts were compared, including Set-1: the original 29,434 SNPs on the 44K chip; Set-2: the 1,090 SNPs overlapped between the 156,502 tagSNPs and 29,434 SNPs; Set-3: the 1,090 SNPs randomly selected from the 29,434 SNPs; Set-4: the 1,090 SNPs evenly distributed in the genome (350 Kb per SNP) selected from the 29,434 SNPs; Set-5: 1,090 SNPs localized within a genomic region from the 29,434 SNPs.

#### The fixedSNP panel

In the crop breeding industry, genotyping cost-per-sample is a top-priority factor, since hundreds to thousands of samples are often genotyped in single day. The data then assists a variety of molecular breeding practices, including genomic selection-assisted phenotype prediction, marker-assisted backcrossing, seed purity or genotype heterozygosity analysis, and subpopulation identification. Cost reduction is usually fulfilled by compiling a highly effective marker panel containing only dozens to hundreds of SNPs that are available for high-throughput genotyping platforms, such as Douglas ArrayTape and LGC Omega-F equipment, using a PCR-based KASP™ genotyping assay. These systems allow users to flexibly combine different numbers of SNPs and DNA samples using multiple plates with 96 and 384 wells per run. To meet the industrial demand, further compression of the tagSNP panel must consider not only the genetic relationship between subpopulations and accessions, but also the evolutionary and/or functional significance of SNPs with high diagnostic effectiveness and stability.

The *Fst* and *θπ* values are commonly used indicators of genomic regions demonstrating signatures of selective sweeps, caused by domestications, artificial selections and environmental adaption. SNPs in selective-sweep regions are usually evolutionarily fixed with strong positive selection signals. To generate the fixedSNP panel, we first identified the selective sweep regions that are specific to each subpopulation and are common to the six subpopulations by combining the ratio of *Fst* versus *θπ* based on the comparison of the cultivated subpopulation against the wild rice population (**Materials and Methods**). Using 100 Kb and 10 Kb windows, large and small genomic regions showing selective sweep signals were identified, respectively. In total, 227 (cultivated *vs.* wild), 381 (*Ind vs.* wild), 333 (*Aus vs.* wild), 296 (*Aro vs*. wild), 256 (*TrJ vs*. wild) and 269 (*TeJ vs*. wild) identified regions showed significantly smaller *Tajima’* D values compared to other regions (**Figure 6A**). Subsequently, genes located in the selective sweep regions and their corresponding GSEA (Gene Set Enrichment Analysis) terms were further identified for each subpopulation, and ∼50% of them were specific to each subpopulation whilst only 27 GSEA terms co-exist in the five cultivated rice subpopulations (**Figure 6B**). Finally, a total of 1,180 SNPs occurred within the genes in the selective sweep regions were selected to generate the fixedSNP panel.

**Figure 6.**
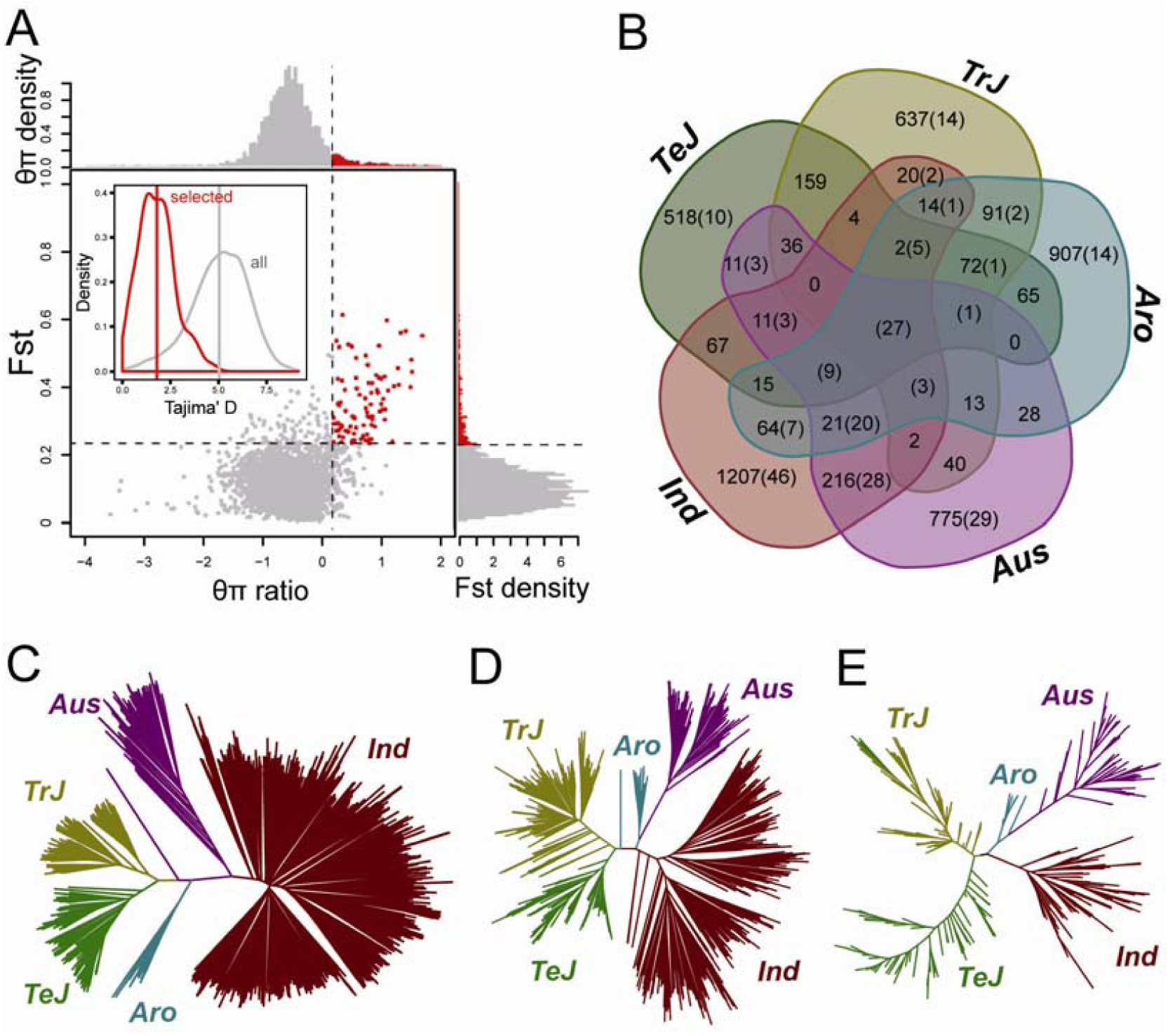
Screening and validation of FixedSNPs. **A**. Distribution of *θπ* ratios (wild vs cultivar) and corresponding *Fst* values, which are calculated in 100kb windows. Data points located to the right of the vertical dashed line and to the top of the horizontal dashed line are potential strong selective sweep signals (Red points, corresponding to the 5% right tails of the empirical *θπ* ratio and *Fst* values distribution, respectively). Distribution of Tajima’s D values for the potential selective sweep signals and whole genomes are shown within the plot. Other comparisons for the screening of subgroup specific selective sweep signals were not shown here, but demonstrate similar trends. **B**. Common and specific selective signals among cultivar subgroups (Number of genes or GSEA terms are shown out and in the brackets, respectively). **C**. Phylogenetic tree of 2,538 rice cultivars in fixedSNPs data set. **D**. Phylogenetic tree of 880 rice cultivars in 700K chip data set. **E**. Phylogenetic tree of 351 rice cultivars in 44K chip data set.

#### Subpopulation classification analysis with the fixedSNP panel

To evaluate the fixedSNP panel, subpopulation classification with phylogenetic tree analysis was performed using the 1,180 fixedSNPs, and the results were compared to the results generated from the 156,502 tagSNPs performed on the same population of 2,556 accessions. All of the accessions were assigned to the correct subpopulations with tagSNPs and the phylogenetic tree showed consistent structure with the tree constructed with fixedSNPs (**Figure 6C**). To further evaluate the universality of the fixedSNP panel, we performed subpopulation classification on two external populations genotyped by SNP chips [11] [14]. One chip dataset contained 880 cultivated rice accessions genotyped by the Affymetrix 700K SNP chip, while the other contained 351 cultivated accessions genotyped by the Illumina 44K SNP chip. Both external chip datasets have been documented with clear subpopulation classification and origins, and possess relatively high genetic diversity. Only 314 and 63 SNPs from the 700K and 44K chips, respectively, were found in the 1,180 fixedSNP panel. For the chip dataset containing 880 accessions, 877 accessions were correctly assigned to their documented subpopulations; three *TeJ* accessions (IRGC121549, IRGC121520 and IRGC121535) were incorrectly assigned to the *TrJ* subpopulation (**Figure 6D**). As for chip dataset containing 351 accessions, 348 were assigned to the correct subpopulation; three *TeJ* accessions (NSFTV134, NSFTV204 and NSFTV283) were mistakenly assigned to *Trj* rice (**Figure 6E**). Overall, 99.8% of the rice accessions examined were assigned to previously documented subpopulation records using markers extracted from the fixedSNP panel, indicating that the fixedSNP panel is an efficient, accurate new tool for subpopulation classification.

#### The barcodeSNP panel

DNA fingerprinting technology using a small set of SNPs to generate a series of genotype combinations, referred to as “barcodes,” has become an economical means to protect commercialized varieties. Thus, the barcodeSNP panel must be able to uniquely identify these barcodes to distinguish between each of the rice varieties on the market. To ensure highest uniqueness but lowest count of barcodeSNPs, we applied the MinimalMarker algorithm on the fixedSNP panel to exhaustively traverse all possible genotype combinations that would distinguish the 2,556 accessions (**Materials and Methods**). The MinimalMarker algorithm generate three sets of minimum marker combinations, in which each set contains 28 SNPs. After merging the three sets, 38 barcodeSNPs were finally selected to generate the panel (**Figure S2A**). In addition, up- and down-stream flanking sequences were also provided for users to design primers for PCR-based KASP™ genotyping assays. The SR4R also offers a web interface that allows users to identify corresponding accessions or varieties when rice varieties are submitted for genotyping with any number of barcodeSNPs between 8 to 38. The SR4R returns a list of the top 10 best-matched accessions/variety in the database, and displays associated information including the accession/variety IDs, number of mismatched bases, genomic position of the barcode, genotype heterozygosity, and documented subpopulation and origin. Among the top 10 hits, if multiple best-matched varieties with 100% identity are returned using a certain number of barcodes, the users may genotype additional barcodeSNPs until a unique best matched variety is identified. It is worth noting that because the SR4R does not have a complete list of the barcodes for all commercial rice varieties in the database, the 38 barcodeSNPs is considered as an initial panel for users to test the best combinations with the most optimal sensitivity and specificity using flexible numbers of markers.

#### Machine learning analysis with the barcodeSNP panel

If a new variety genotyped with barcodeSNPs is not found in the database, SR4R will perform subpopulation classification. The traditional method of subpopulation classification first integrates the genotype of the submitted variety with the genotypes of all the varieties in the database, then performs phylogenetic analyses to determine the best assigned subpopulation. This procedure is tedious and computationally inefficient since the database contains hundreds of thousands of accessions. To simplify the procedure so that it may be implemented through a web interface, we adopted an alternative method that utilizes machine learning-based subpopulation classification models with the 38 barcodeSNPs as features. We used all of the 2,556 rice accessions to evaluated seven commonly used machine learning algorithms to perform subpopulation classification including decision tree, k-nearest neighbouring, naïve Bayesian, artificial neural network, random forest, multinomial logistic regression and one-*vs*-rest logistic regression algorithms, followed by ten-fold cross validation assessment (**Materials and Methods**). A series of assessments of the classification precision in the five cultivated rice subpopulations indicated that, out of the seven methods the best one is the multinomial logistic regression model, whose AUC (Area under curve) values were all ≥ 0.99 for all subpopulations (**Figure S3B-F**). Additional methods are one-vs-rest logistic regression and the random forest model; where results from each yielded similar classification precision to the multinomial logistic regression model. Then, we used an independent datasets containing 880 rice accessions profiled by 770 Kb rice SNP chip for independent validation. The multinomial logistic regression model was trained by the 2,556 rice accessions, and then predict the subpopulation classifications on the 880 samples. The AUC values were all ≥ 0.99 for all subpopulations in this independent datasets, indicating robustness of the model. Moreover, compared the original label and the predicted label with the max probability for each sample, the true positive rate (TPR) and false positive rete (FPR) are also reasonable (**Figure S4**). The pre-trained classification models with the seven machine learning algorithms have been implemented on the SR4R server provided as a web tool for users to perform subpopulation classification when the genotype information of the 38 barcodeSNPs in submitted.

#### The barcodeInDel panel

InDel (Insertion and Deletion) is another form of genomic variations (usually less than 50 bp in length) that can be used as molecular markers for a variety of population analysis. From the 5,152 rice accessions, a total of 4,217,174 raw InDel variations were identified using the IC4R variation calling pipeline [2]. After filtering low-quality InDels, 109,898 high-confidence InDels were retained with missing rate less than 0.01 and MAF ≥ 0.05 within 2,556 rice accessions. Among the 109,898 high-confidence InDels, we further identified 62 subpopulation-specific InDels which can be used as barcodeInDels to differentiate the six rice subpopulation *TeJ, TrJ, Aro, Aus, Ind* and *Oru*, and the six subgroups of *indica* rice *S1-S6* (**Table S3**). The 109,898 high-confidence InDels can be download from SR4R for users’ customized analysis.

#### Web interface of SR4R database

Using unified bioinformatics pipelines, the genotype data of the 18 million raw SNPs identified from 5,152 rice accessions were processed to construct four reference panels of SNPs for different utilizations. Because genotype data processing is a complicated and computationally intensive procedure, the four SNP panels are readily usable for a variety of analyses simplify task for rice researchers. For better sharing of SNPs and improvement of the rice variation map utility, we developed the SnpReady for Rice (SR4R) database. Through the SR4R web interface, users may directly browse the four panels and retrieve detailed information related to the 2,097,405 hapmapSNPs, 125,502 tagSNPs, 1,180 fixedSNPs, 38 barcodeSNPs. In addition, the protein-coding genes exhibiting strong selection signatures, associated with the 1,180 fixedSNPs were also included in the SR4R database with detailed functional annotations (**Figure 7A**). When users retrieve a SNP such as the first SNP “OSA01S00001362”, the genomic location and the adjacent gene or the gene containing the queried SNP are displayed. Users may also retrieve a visualized allele frequency map in the six major subpopulations, and the six subgroups of *indica* rice (**Figure 7B**).

**Figure 7.**
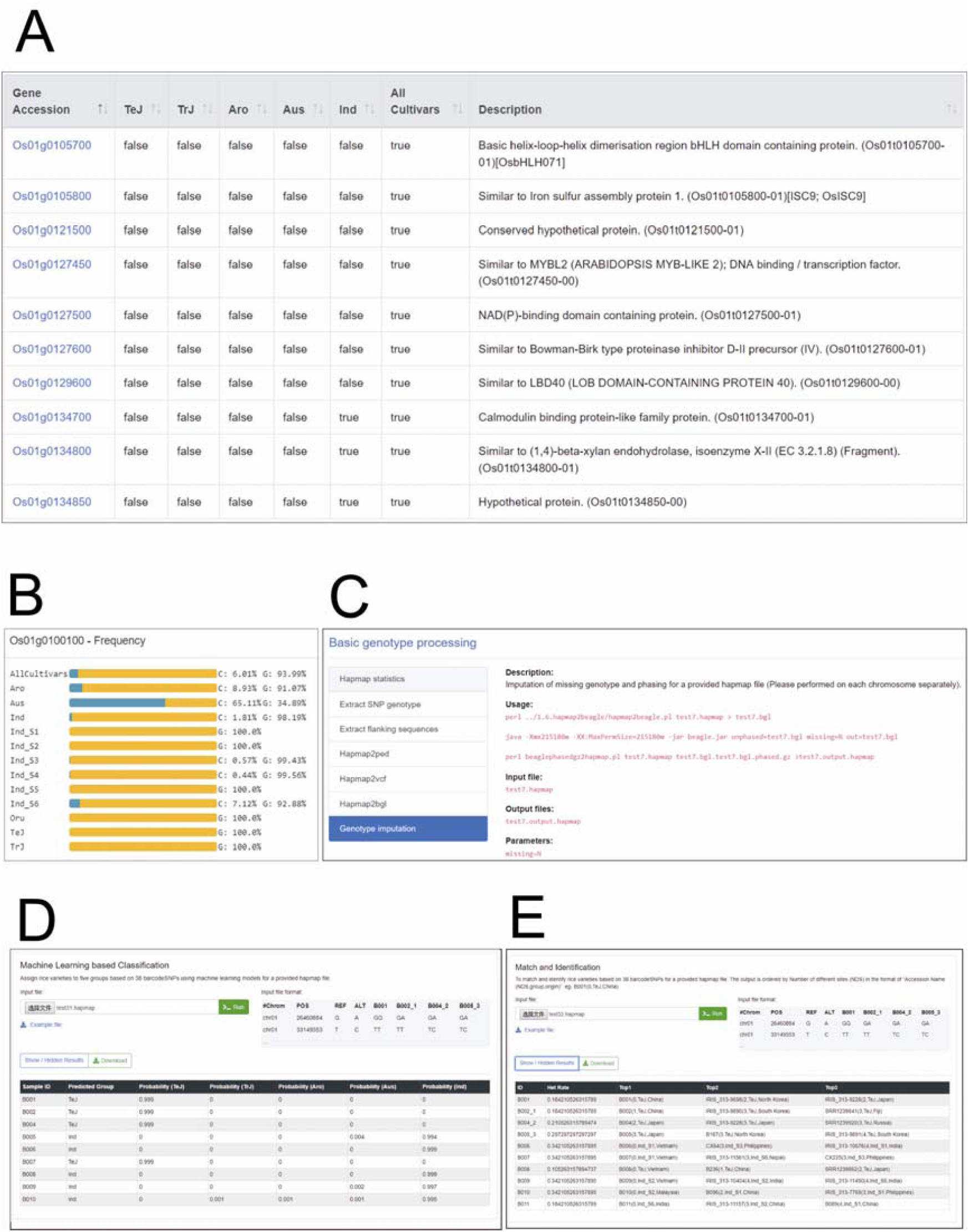
Representative functional modules in SR4R database. **A.** Genes exhibiting significant selection signatures in the corresponding subpopulations are listed in the “Selected Genes” module in the Browser. **B.** Allele frequencies in different subpopulations of the first hapmapSNP (SNPID: OSA01S00001362, associated gene: Os01g0100100, position: chr01-1362, allele: Alt-A, Ref-G). **C**. One example of the script and pipeline for population diversity analysis. **D**. The online analysis module of subpopulation classification using machine learning algorithms. **E**. The online analysis module of rice variety identification using the 38 barcodeSNPs.

The users may also download the four panels of SNPs along with the original genotype files for local analysis *via* http://sr4r.ic4r.org/download. In addition, the “Tools” module presents 18 handy scripts and pipelines that users may install on their local computers for a variety of analysis, including basic genotype processing, population diversity analysis, rice variety identification and subpopulation classification. For example, assuming one user may want to perform a genotype imputation of 44K SNP rice Chip, she or he may first download the file “hapmapSNPs-genotype.tar.gz (892 MB)” containing the genotypes of the 2,097,405 hapmapSNPs in 2,516 rice accessions. Then, the user may use the pipeline and scripts demonstrated in **Figure 7C** to perform imputation on a local server. SR4R also offers two modules of online analysis. The first module is to use a machine learning-based method to assign the subpopulation type based on the user-submitted genotype file including no more than 20 samples. The model will return the probability of the type of subpopulation assigned to each sample (**Figure 7D**). The second module is to perform DNA fingerprint analysis. When the user submits a genotype file containing no more than 20 samples, the model will search the accession database, and return the top three matches of existing varieties with the number of mismatched nucleotide and heterozygosity rate displayed (**Figure 7E**). The programs and scripts for these two modules along with demo input and output files are also available to download for local analysis of genotypes with large sample numbers.

## Conclusions

The IC4R Rice Variation Database collects over 18 million raw SNPs identified from resequencing of 5,152 accessions. To meet the different demands for the rice research community and breeding industry, we further generated four panels of 2,097,405 hapmapSNPs, 156,502 tagSNPs, 1,180 fixedSNPs and 38 barcodeSNPs with standard processing pipelines and uniform analytical parameters (**Table S2**). The four panels of SNPs can be either accessed online or downloaded for local use from the daughter database of RVD – SnpReady for rice (SR4R). The hapmapSNP panel contains 2 million non-missing genotypes of 2,556 accessions offers a reference HapMap for genotype imputation and high-resolution GWAS analysis. The non-redundant 150K tagSNP panel is an ideal magnitude for population genetics and evolutionary analysis for research, as well as an ideal marker pool for genomic selection-assisted breeding in rice. For a breeding population with about 500 F_1_ hybrids, 1,500 to 15,000 markers selected from the tagSNP panel can be used to build a GS model, reaching a satisfactory genotype-to-phenotype prediction accuracy. The fixedSNP panel with high effectiveness and stability can be regarded as a marker pool for various molecular breeding practice suitable for low-budget, flexible genotyping platform, in terms of subpopulation classification, seed purity analysis and genetic background analysis. The 38 barcodeSNPs selected by MinimalMarker algorithm is an initial marker set for generating DNA fingerprints for commercial rice varieties. Along with the barcodeSNP panel, two web-based tools, one for variety identification and another for subpopulation classification, are offered in SR4R. In addition, the SR4R database also offers a series of standard pipelines used to construct the four sets of SNPs, and local handy tools to perform rice varieties classification, barcode development, and other types of genetic and breeding research. With the incremental accumulation of population genotype data in BIGD center, these bioinformatics tools can be applied to other animal or plant species such as corn, wheat, soybeans, for a centralized reference HapMap and SNP panel databases for plants.

## Materials and Methods

### Construction of hapmapSNP and tagSNP panels

The raw 18 million SNPs with genotype information of 5,152 rice accessions were obtained from the IC4R rice variation database (http://variation.ic4r.org). Accession filtration, SNP filtration and basic statistics of homozygous SNPs and accession heterozygosity were performed using in-house scripts. Genotype imputation of missing sites and phasing were performed using Beagle [8]. A SNP site with missing genotype was removed if an inferred genotype with a posterior probability was smaller than 0.5. Genomic annotation of hapmapSNPs was performed using ANNOVAR (version 20160201) against the rice International Rice Genome Sequencing Project (IRGSP) gene annotation. Using the reported LD length of rice ranging from 40 to 500 Kb, an LD-based SNP pruning method was used to construct the tagSNPs category using PLINK with –*indep* command [15] [16]. The PLINK parameters were selected based on the variance inflation factor (VIF), which recursively removed SNPs within a sliding window of 50 SNPs and a step size of 5 SNPs to shift the window.

### Tools for subpopulation structure analysis

The tagSNPs for 2,556 rice accessions were concatenated as input sequences for constructing the phylogenetic tree using the neighbour joining algorithm implemented in MegaCC with pairwise gap deletion and 100 bootstrap replications [17]. The output tree file for all 2,556 rice accessions and the subtree file of *indica* rice accessions were visualized in MEGA7 [18]. Principal component analysis of the 2,556 rice accessions was done by flashPCA [9]. Population admixture structure analysis was done by fastSTRUCTURE using the variational bayesian framework, and *k=2* to *k=8* were set to infer the admixture of ancestors for the accessions.

### Tools for genetic diversity analysis

Genetic diversity related analyses were mostly done using PLINK [16]. Genome-wide pairwise IBS calculations were performed between each pair of accessions within the same subpopulation in order to deduce the genetic affinity, and an IBS pairwise distance matrix was generated for each subpopulation. The ROH analysis for each subpopulation used a sliding window method to scan each accession’s genotype for a given population at each marker position to detect homozygous segments. The parameters and thresholds applied to define ROH were set as follows: a minimum ROH length of 200 Kbp and a minimum number of 1,000 consecutive SNPs included in an ROH. Correlation coefficient (*r*^2^) of SNPs was calculated to measure LD level for each subpopulation. The average *r*^2^ value was calculated for each length of distance from 0 to 500 Kbp, followed by drawing LD decay figures using an R script for each subpopulation. Population diversity of rice varieties was measured by two indexes: *θπ* and *Fst.* Nucleotide diversity *θπ* was used as a measurement of the degree of genotype variability within each subpopulation, while subpopulation differentiations were evaluated by the fixation index *Fst* for each of the cultivated subpopulations against the wild rice population and for the cultivated subpopulations compared to each other. Values of *θπ* and *Fst* were calculated using sites mode implemented in VCFtools [19].

### Tools for genomic selection analysis

Genotype and phenotype datasets of the 44k rice chip were downloaded from the Rice Diversity Website (http://www.ricediversity.org/). Genotype imputation and phasing were then performed using Beagle (version 3.3.2), and the site was filtered if an inferred genotype with a posterior probability was smaller than 0.5. Genomic selection analysis was performed using RR-BLUP mixed model implemented in R package rrBLUP [13] for nine well-measured traits (flowering time, panicle fertility, seed width, seed volume, seed surface area, plant height, flag leaf length, flag leaf width, and florets per panicle) with five different feature combinations. The prediction accuracy under each feature combination was evaluated by five-fold cross-validation and Pearson correlation coefficient. An example of the process is as follows: the original samples were randomly partitioned into five subsets; of the five subsets, a single subset was retained as the validation data, and the remaining four subsets were used as training data. This process was repeated five times, with each of the ten subsets used exactly once as the validation data. The Pearson correlation coefficients of the predicted breeding values and the real phenotype values were calculated for each fold.

### Construction of the fixedSNP panel

*θπ* and *Tajima’* D values were calculated for six rice subpopulations (*TeJ, TrJ, aro, aus, ind, Oru*) with a sliding-window fashion across the genome using in-house scripts. *Fst* values were calculated for the five cultivated subpopulations against the wild *Oru* subpopulation, as well as for the five cultivated subpopulations against each other. For each pairwise comparison, the intersection of the top 5% windowed *θπ* ratios (wild subpopulation *vs*. cultivated subpopulation), and the top 5% windowed *Fst* values correspondingly were selected as strong selective sweep signals. Window sizes of both 100 Kbp and 10 Kbp were used to detect large or small selective sweep regions, followed by merging the results as the candidate selective sweep regions for each subpopulation. *Tajima’* D distribution was also drawn for the candidate selective sweep regions against the whole genomes for each pairwise comparison. Genes located within the candidate selective sweep regions were extracted for each comparison, and Gene Set Enrichment Analysis (GSEA) was performed for each gene listed by using PlantGSEA web tools [20]. Genic SNPs located in the candidate selective sweep regions identified from the above-mentioned pairwise comparisons were merged as fixedSNPs.

### Construction of the barcodeSNP panel

The 1,180 fixedSNPs were used as the initial marker set to select the minimal number of barcodeSNPs that can maximally distinguish the 2,556 rice accessions using a heuristic mode implemented in MinimalMarker [7]. Three minimal sets each containing 28 SNPs were generated, and after merging the three sets, 38 unique SNPs were selected as barcodeSNPs for generating DNA fingerprints for each accessions.

To identify commercialized rice varieties using the combination of 38 barcodeSNPs, seven machine learning-based methods were used: decision tree, k-nearest neighboring, naïve Bayesian, artificial neural network, random forest, multinomial logistic regression, and one-vs-rest logistic regression algorithms in the Python sklearn library (https://scikit-learn.org/stable/). The precision of each model was assessed using ten-fold cross-validation method. Specifically, the original sample set was randomly partitioned into ten subsets in which nine subsets were used for training model and the remaining subset was used as the testing model; this procedure was repeated ten times and an average prediction accuracy was computed from the overall performance of the tested models. Five one-hot codes (10000, 01000, 00100, 00010, 00001) to label the five subpopulations for classification using machine learning models. Then, the predicted label with the max probability was compared with the original label for each sample. If the predicted label is identical with the original label, the prediction result was regarded as correct. Then, the ratios of positive and negative rate were computed to plot ROC curves and compute AUC values.

### Construction of the barcodeInDel panel

Raw InDels were identified using the IC4R variation calling pipeline from the origin 5,152 rice accessions [2]. Then, the InDels from the 2,556 rice accessions with high sequencing coverage (depth ≥ 5) presented in SR4R database were extracted using customized Python scripts, followed by using VCFtools [19] to filter InDels to generate a high-confidence InDel dataset, with parameters of missing rate less than 0.01 and MAF ≥ 0.05. Finally, using customized Python scripts, InDels which have the same sequence type within each subpopulation were retained to generate the subpopulation-specific barcodeInDel panel.

## Data availability

All the data is freely available and downloadable at http://sr4r.ic4r.org/.

## Authors’ contributions

XFW, SHS and ZZ conceived the project; JY and CL collected the samples; JY conducted the data analysis; DZ developed the database; JY, XFW, SHS and ZZ wrote the manuscript.

## Acknowledgments

We are grateful to a number of users for reporting bugs and providing suggestions in improving SR4R. This work was supported by the National Science Foundation of China [31871706], by the Department of Agriculture of Guangdong Province (2018-36), Science and technology program of Guangdong Province (2019B030316006) and by The Youth Innovation Promotion Association of the Chinese Academy of Sciences [2017141].

## Supplementary material

**Figure S1.**
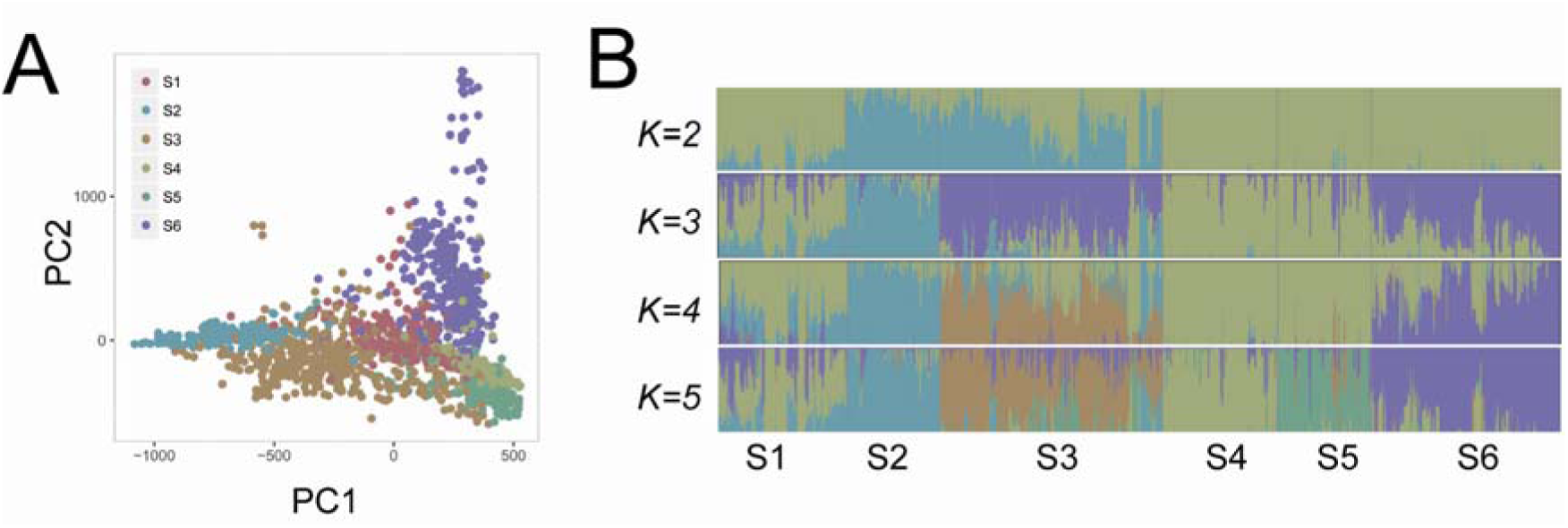
Population structure of 1,655 varieties in *indica*. **A**. PCA classification for 1,655 varieties in *indica* subgroup. **B**. Structure analysis for 1,655 varieties in *indica* subgroup.

**Figure S2.**
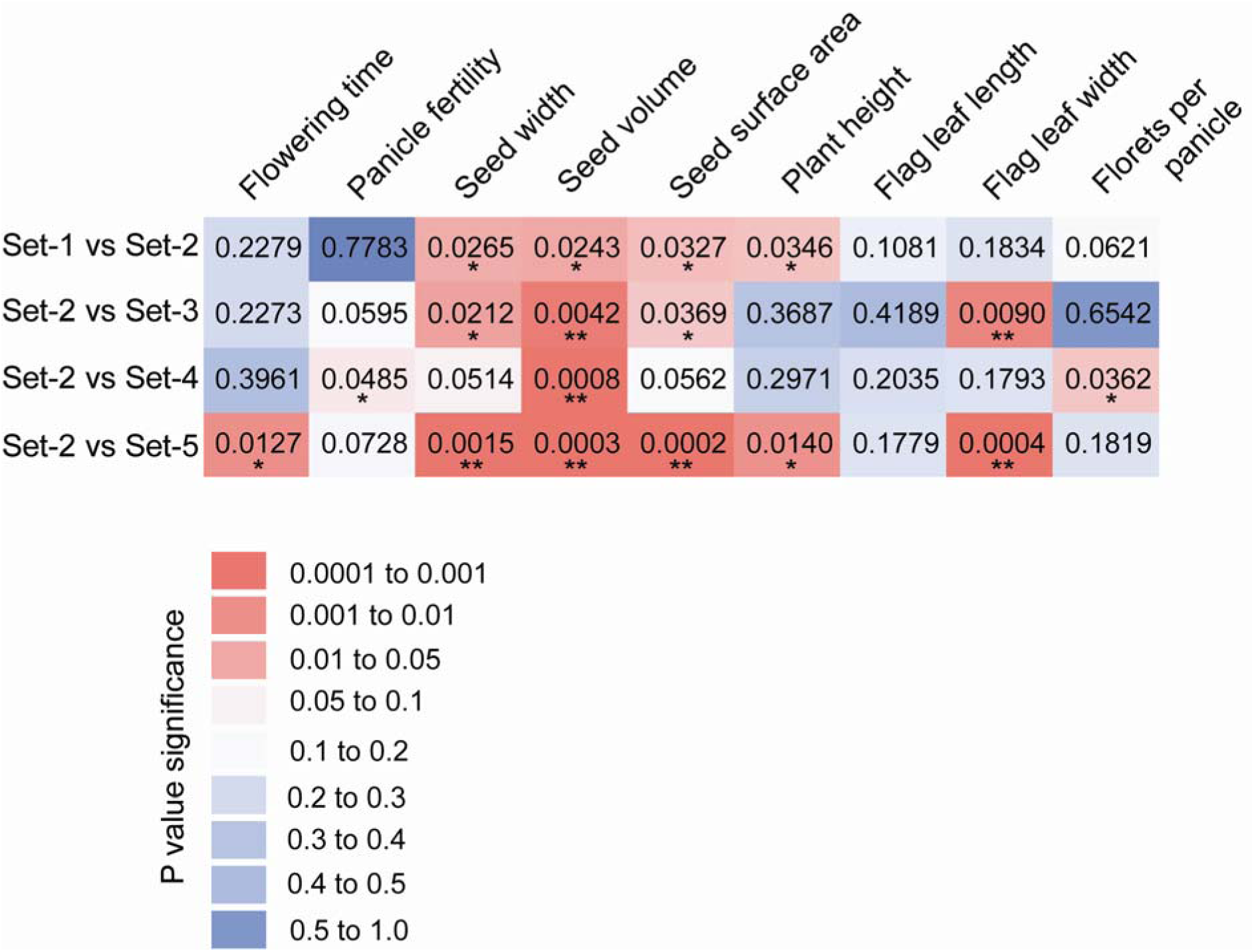
T-test for Pearson correlations of the selected 1,090 tagSNPs set and other four SNP sets. Set-1: the original 29,434 SNPs on the 44K chip; Set-2: the 1,090 SNPs overlapped between the 156,502 tagSNPs and 29,434 SNPs; Set-3: the 1,090 SNPs randomly selected from the 29,434 SNPs; Set-4: the 1,090 SNPs evenly distributed in the genome (350 Kb per SNP) selected from the 29,434 SNPs; Set-5: 1,090 SNPs localized within a genomic region from the 29,434 SNPs. Different colors present different P values. * P value < 0.05; ** P value < 0.01.

**Figure S3.**
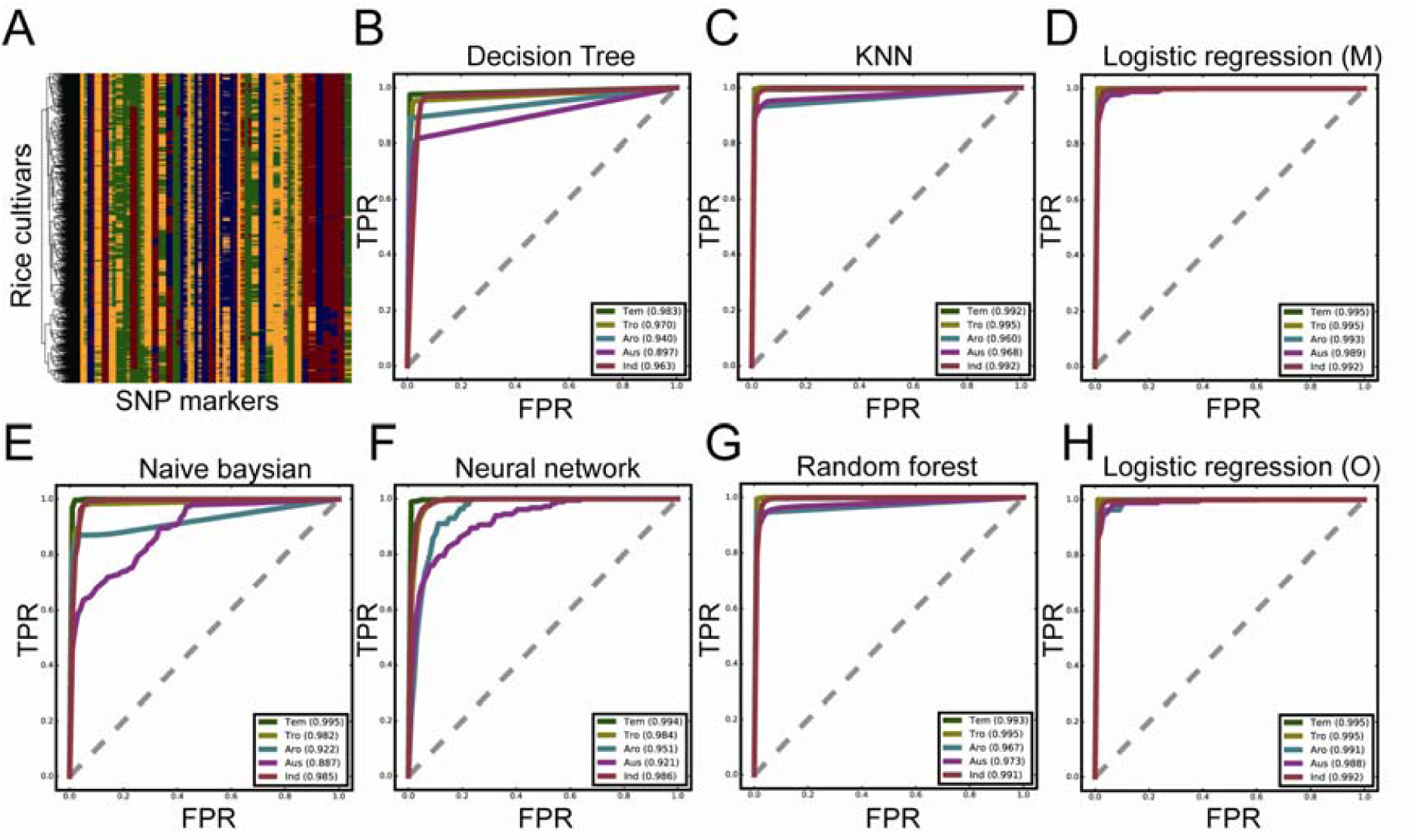
BarcodeSNPs and machine learning models for the classification of rice varieties. **A.** Heat-map of BarcodeSNPs of 2,538 rice cultivars (Red: A, Yellow: T, Blue: G, Green: C). **B.** Decision model. **C**. KNN model. **D**. Multinomial logistic regression model. **E**. Naive Bayesian model **F**. Neural network model. **G**. Random forest model. **H**. One-vs-rest logistic regression model. AUC curves were drawn using the mean values of ten cross validations for B-H.

**Figure S4.**
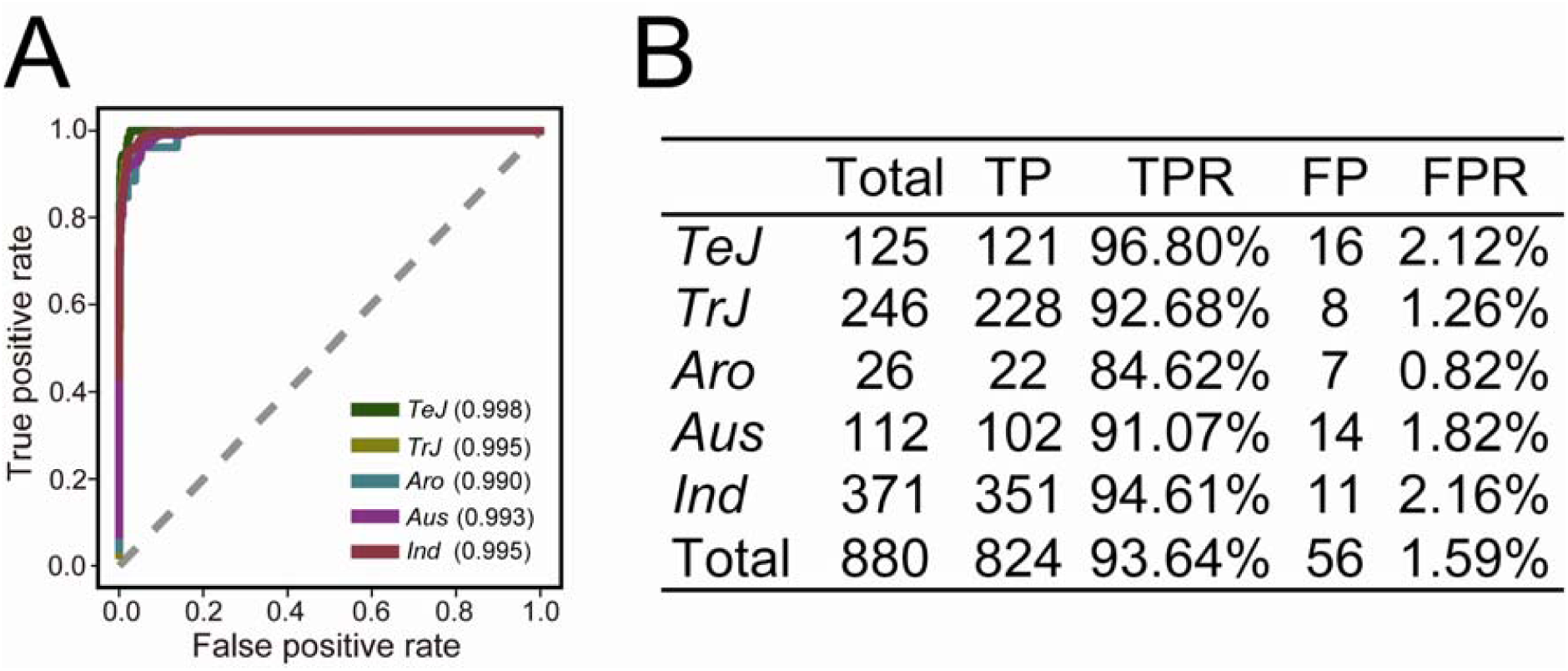
Independent validation of the machine learning model. **A**. ROC curve for the 770 Kb rice SNP chip dataset using the pre-build multinomial logistic regression model. **B.** The true positive rate (TPR) and false positive rate (FPR) statistics for each subpopulation of the 770 Kb rice SNP chip dataset.

**Table S1 Summary of 2,556 rice accessions with subpopulation classification and origins**

**Table S2 Summary of SNPs annotation for SR4R database**

**Table S3 The barcodeInDel panel in SR4R database**

